# Subdiffusive random growth of bacteria

**DOI:** 10.64898/2026.03.19.712816

**Authors:** Jialiang Wei, Yichen Yan, Jie Lin

## Abstract

While the regulation of bacterial cell size is widely studied across generations, the stochastic nature of cell volume growth remains elusive within a cell cycle. Here, we investigate the fluctuations of cell volume growth and report a subdiffusive random growth. Specifically, the mean square displacement of the logarithmic volume scales as Δ*t*^*α*^ with an anomalous exponent *α* ≈ 0.27. This low exponent implies strong negative temporal correlations in growth rate noise on timescales of minutes, which are significantly faster than those of gene expression dynamics. We attribute this phenomenon to the physical mechanics of the cell wall. By modeling the cell wall as a complex viscoelastic material with power-law-distributed relaxation times, we successfully recapitulate the observed subdiffusive behavior. Our results suggest that it is the heterogeneous cell-wall viscoelasticity, rather than biological regulatory programs, that governs the short-timescale fluctuations of bacterial growth.

## Introduction

Understanding how bacteria regulate their size is a fundamental question in quantitative biology [1, 2]. Phenomenological models, such as the “adder” model, have successfully described cell size control at the intergenerational level [3–5]. Regarding the average growth dynamics within a cell cycle, it has been suggested that bacterial cell volumes grow exponentially over time [4–7]. Nevertheless, recent studies on *Escherichia coli* and *Caulobacter crescentus* have suggested that growth is not strictly exponential but exhibits a “superexponential” acceleration toward the end of the cell cycle [8–10]. However, it remains unclear whether the observed growth rate variations reflect intrinsic biological programs or arise as artifacts of the statistical methods used to analyze stochastic trajectories. What is even more elusive is the stochastic nature of cell volume growth on top of the deterministic growth, and how a bacterial cell regulates its volume on a shorter time scale than the cell cycle.

In this Letter, we employ a minimal drifted Brownian model as a controlled benchmark to evaluate common statistical frameworks. We demonstrate that conditioning average growth rates on relative age, volume, or elapsed time since cell birth inherently introduces systematic biases that can be mistakenly interpreted as biological signals. After accounting for these artifacts, our analysis of experimental data of *E. coli* suggests a genuine growth rate acceleration in the end of the cell cycle, supporting the conclusion of superexponential growth.

More significantly, we investigate the fluctuations of cell volume growth on top of the non-exponential deterministic growth. Strikingly, the random growth rate of cell volume exhibits a subdiffusive fluctuation. Suppose we consider the growth rate as the velocity of an imaginary particle and the logarithmic volume 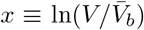 as the particle position, the mean square displacement (MSD) of *x* scales as 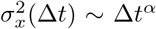 with an anomalous exponent *α* ≈ 0.27. This subdiffusion implies that the growth rate noise exhibits negative temporal correlations. Given that the subdiffusion occurs on a time scale of minutes, much shorter than the time scales of significant changes in protein concentrations [11, 12], we hypothesize that this phenomenon has a physical origin.

Given the fact that the deformation of the cell wall is a prerequisite of cell volume growth, we propose that the subdiffusive random growth of cell volume is a consequence of the mechanical properties of the cell wall [13–17]. Specifically, we model the cell wall as a complex viscoelastic material composed of multiple serially connected Voigt elements with power-law-distributed relaxation times. This simple model successfully recapitulates subdiffusive growth behavior, and our results suggest that the viscoelastic and heterogeneous cell wall fundamentally constrains bacterial growth.

### Cell cycle-dependent growth rate

We introduce a minimal model of cell volume growth as a control for the experimental data and define 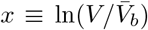 where *V* is the cell volume and 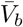 is the birth volume averaged over cell cycles, which is a constant. In the minimal model, *x* follows a drifted Brownian motion:

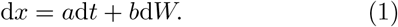

Here, *t* is time, *a* is the deterministic growth rate, and *b* is the noise intensity. d*W* represents a Wiener process increment, which is a Gaussian-distributed random number with mean equal to zero and variance equal to d*t* [18]. Cell division is implemented using the adder model [1, 3, 5]: *V*_*d*_(*n*) = *V*_*b*_(*n*) + *V*_0_ + *ζ* and *V*_*b*_(*n* + 1) = *V*_*d*_(*n*)*/*2. Here, *n* is the generation number, *V*_*d*_ is the division volume, *V*_*b*_ is the birth volume, *V*_0_ is a constant, and *ζ* is a Gaussian-distributed random variable. The simulated cell volume trajectories resemble the experimental ones (Fig. 1(a, b); see simulation details in End Matter).

**FIG. 1.**
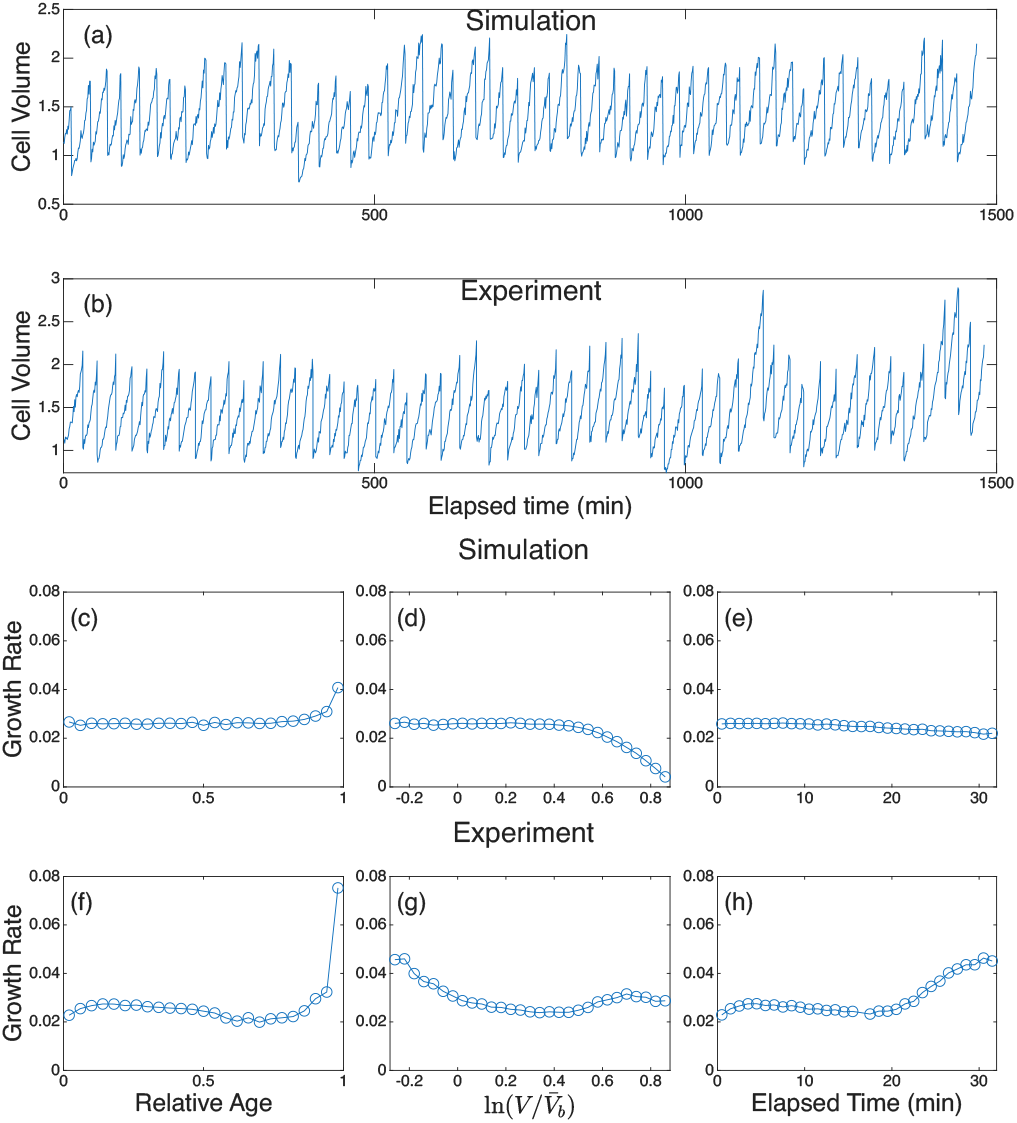
Cell cycle-dependent average growth rate. (a) Cell volume vs. time for the simulation data, where the cell volume is normalized by the average birth volume 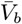. Here, *a* = 0.026, *b* = 0.040 with the time unit 1 min. (b) The same as (a) but for the experimental data. (c-e) The growth rate *λ* = ⟨Δ*x/*Δ*t*⟩ averaged over cell cycles for the simulation data vs. relative age (c), 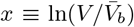 (d), and the elapsed time (e). (f-h) The same analysis as (c-e) but for the experimental data of *E. coli* in the glucose + 12 a.a. medium in Ref. [5].

Previous works have shown that the growth rate *a* was not constant during the cell cycle for *E. coli* and other bacteria [8–10]. To test whether the cell cycle-dependent growth rate is due to real biological processes or data analysis artifacts, we compute the growth rate as *λ* = ⟨Δ*x/*Δ*t*⟩ using the same time interval (Δ*t* ≈ 1 min) across both experimental and simulation datasets. Because the growth rate is strictly a constant *a* in the minimal model, any deviation of the growth rate from *a* must be a numerical artifact.

We apply three statistical methods to analyze both experimental and simulation data (Fig. 1(c-h); End Matter). Surprisingly, all three statistical methods yield systematic biases (Fig. 1(c-e)). In the relative-age statistics (Fig. 1(c)), we compute the average growth rate conditioned on a given relative age *ϕ* during the cell cycle, which is normalized from 0 to 1. Intriguingly, even the simulation data exhibits an apparent acceleration at the end of the cell cycle. In particular, we prove that the average growth rate diverges as *λ* ∼ (1 − *ϕ*)^−1*/*2^ near the end of the cell cycle, a purely statistical effect (End Matter).

In contrast, in the relative-volume statistics, where we compute the average growth rate conditioned on a given 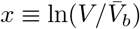, the average growth rate decreases near the end of the cell cycle in the simulation data (Fig. 1(d)). The reduction decreases as the time interval to compute the average growth rate Δ*t* decreases (Fig. S3(b)). A similar process occurs in the elapsed-time statistics, where we compute the average growth rate conditioned on a given elapsed time since cell birth (Fig. 1(e)), Fig. S3(c)). We leave the discussion of the mechanisms underlying these statistical biases for End Matter, which is not the primary focus of this work.

Given the minimal model as a control, our analysis of experimental data supports the conclusion of superexponential growth at the end of the cell cycle, previously proposed in the literature [8–10]. For the relative-volume and elapsed-time statistics, we still observe a growth rate acceleration at the end of the cell cycle (Fig. 1(g, h)), despite the growth rate reduction predicted from the minimal model. We also observe a growth rate acceleration at the end of the cell cycle in the relative-age statistics (Fig. 1(f)); however, this is not irrefutable evidence for a true superexponential growth (End Matter). As we show later, the growth rate noise in experimental data is colored with negative temporal correlation. Nevertheless, the observed growth rate biases for the three statistics are still valid for colored noise (Fig. S6).

### Subdiffusive random growth

Bacterial cell volume is well regulated across cell cycles; however, it is still elusive whether cell volume growth is regulated on a shorter time scale. Therefore, we study fluctuations in cell volume growth using the minimal model as a control. We calculate 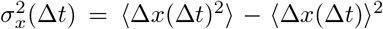 where Δ*x*(Δ*t*) = *x*(*t*_0_ + Δ*t*) − *x*(*t*_0_) given a time interval Δ*t*; here, the average is over cell cycles conditioned on a given relative age, which we call the mean square displacement (MSD) of *x* in the following. For the minimal model, *x* is a random walk with a constant drift; therefore, 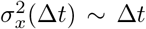 (Fig. 2(a-c)). Indeed, the measured exponent was close to one as long as the relative age is not close to zero or one (Fig. 2(b, c)), from which we introduce a regime where the measured exponent is reliable (the shaded area in Fig. 2(c)). Strikingly, we find that the experimental data of *x* implements a subdiffusive noise (Fig. 2(d-f)):

**FIG. 2.**
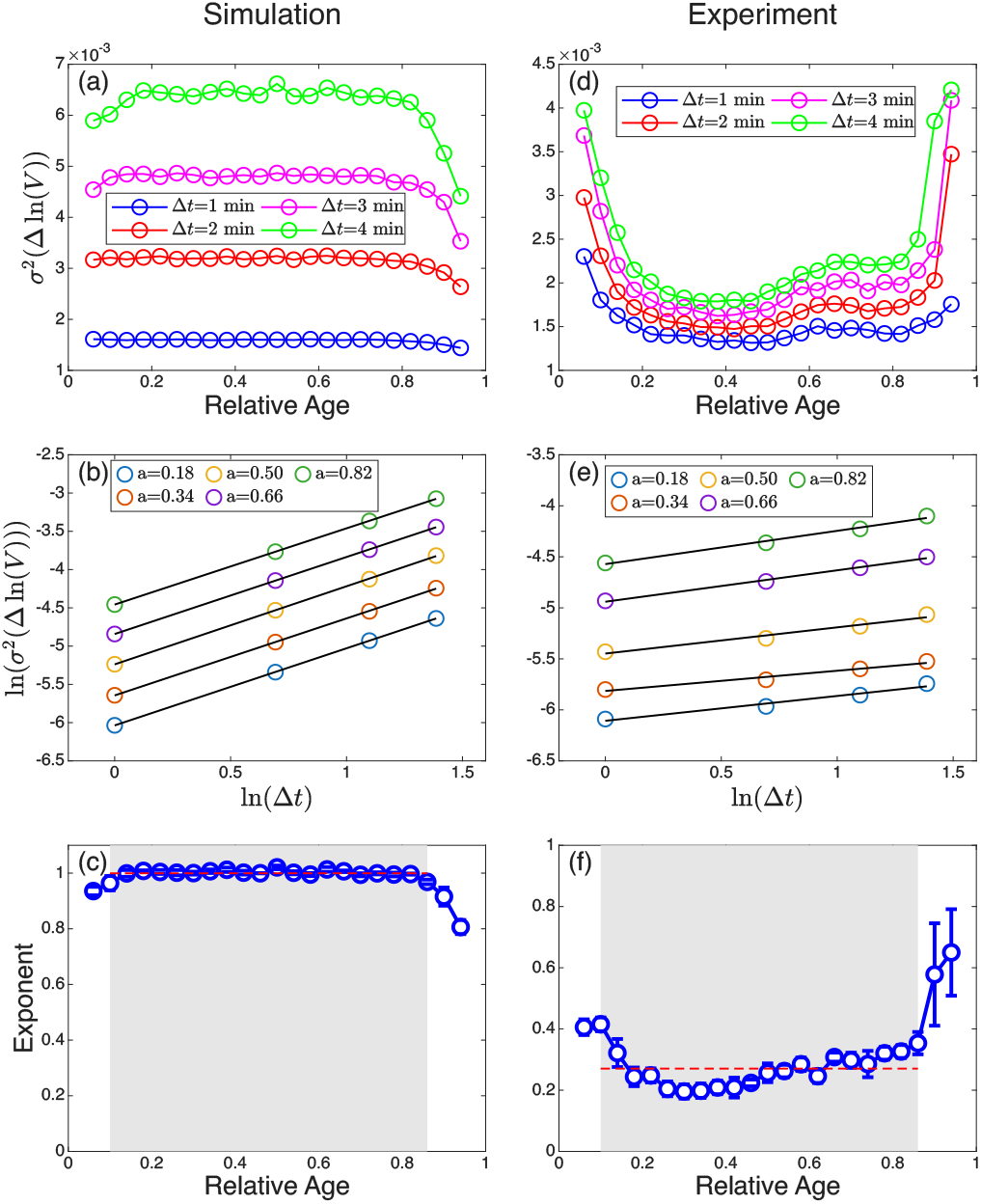
Subdiffusive random growth of cell volume. (a) 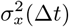 vs. the relative age for time intervals Δ*t* = 1, 2, 3, and 4 min for the simulation data. (b) 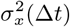 vs. Δ*t* for several relative ages, from which we calculated the diffusion exponent. Curves for different relative ages are vertically shifted for visual clarity to highlight the robust power-law scaling. (c) The diffusion exponent (*α*) calculated from the simulation data, plotted against the relative age, showing the reliable regime where *α* is equal to the ground truth 1. (d-f) The same analysis as (a-c) but for the same experimental data of *E. coli* in Fig. 1. The red dashed line in (c, f) is the average exponent in the reliable regime.

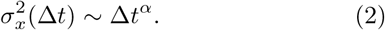

Notably, 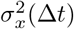 in the experimental data is significantly larger at the early and late stages of the cell cycle (Fig. 2(d)), which may be generated by the morphological rearrangements during cell birth and division. Intriguingly, the *α* exponent changes mildly throughout the cell cycle, with an average value, *α* ≈ 0.27 in the reliable regime. We also analyze the MSD using growth rate data derived from cell length and surface area and find similar subdiffusive behavior (Fig. S4). More importantly, this anomalous scaling is robust across a wide range of growth rates and nutrient conditions (Fig. S5). The subdiffusion suggests that the growth rate noise at different times must be negatively correlated.

### Modeling cell-volume dynamics as a complex viscoelastic material

Because the subdiffusion occurs in a time scale of minutes for the experimental data (Fig. 2(e)), it is unlikely to be generated by biological activities associated with gene expression, which usually takes a doubling time for the protein concentrations to change significantly in bacteria [11, 12]. Therefore, we hypothesize that the subdiffusion comes from a physical process. Considering that cell volume growth always accompanies cell wall growth, and that the cell wall is a complex viscoelastic material [15, 17], we propose that the cell wall mediates the subdiffusive random growth.

We introduce a minimal model of cell-wall mechanics in which we model the cell wall, a peptidoglycan network, as a serial connection of *N* Voigt elements with elastic and viscous components, which we denote as the general Voigt model (Fig. 3(a)). For one particular Voigt element *i*, we define its strain as the deviation from its deterministic growth Δ*ξ*_*i*_ = Δ*x*_*i*_ − *a*_*i*_Δ*t*. For simplicity, we set the growth rate *a*_*i*_ as a constant in the following simulations, which do not affect our conclusions regarding the growth rate noise. The stress *p*, which drives cell wall deformation and cell volume growth, corresponds to the turgor pressure (i.e., the difference between cytoplasmic and external osmotic pressures) shared by all elements [19–23]. Here, because we focus on the growth rate noise, what matters to us is the stress fluctuation, *δp*, which we set as a white noise, such that ⟨*δp*(*t*)⟩ = 0 and ⟨*δp*(*t*_1_)*δp*(*t*_2_)⟩ = 2*A*_*p*_*δ*(*t*_2_ − *t*_1_) where *A*_*p*_ is a constant.

**FIG. 3.**
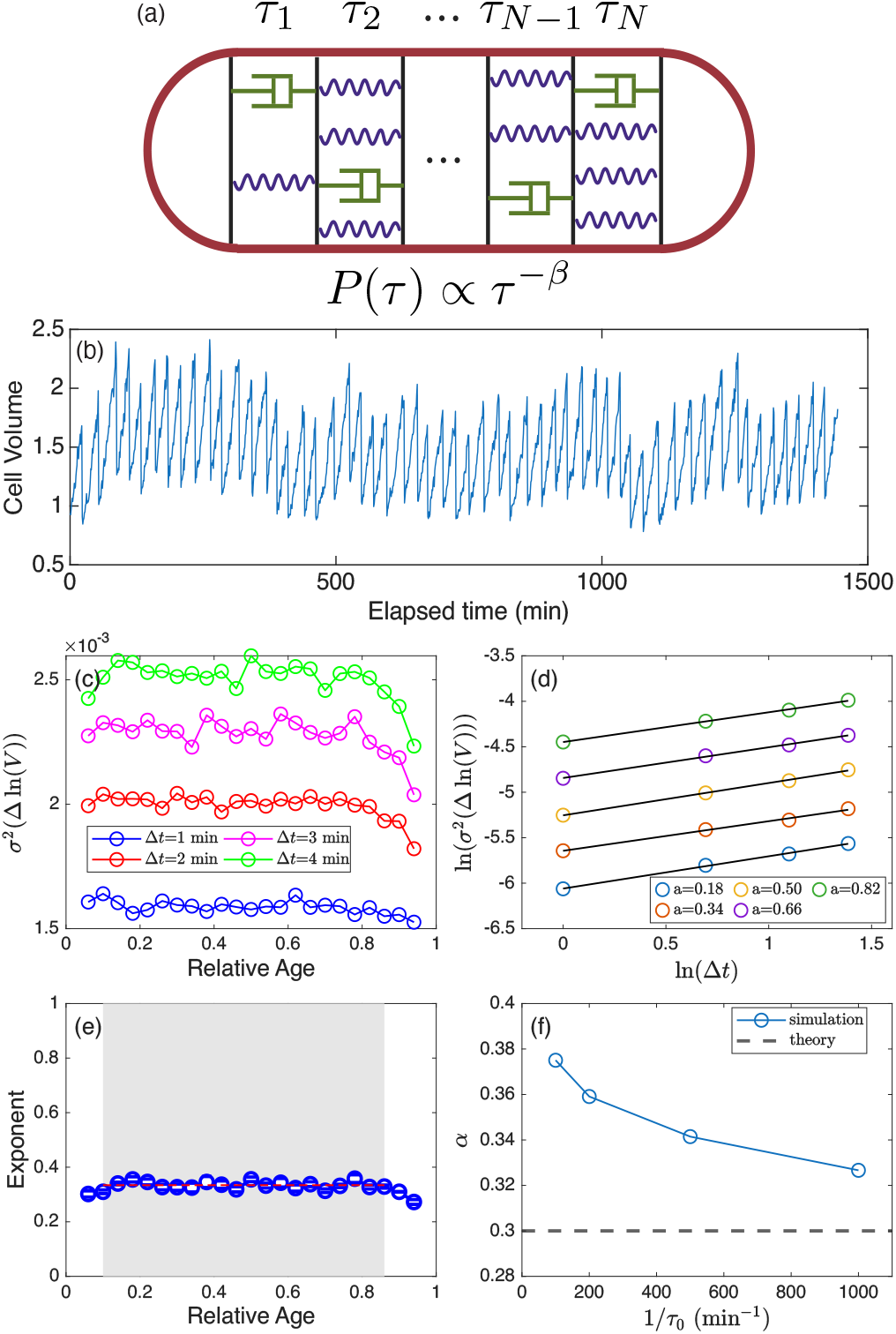
The general Voigt model of the cell wall recapitulates the subdiffusive random growth. (a) We model the cell wall as a serial connection of multiple Voigt elements, each with an elastic component (the spring) and a viscous component (the dashpot). We assume that the relaxation time of each element is random due to the random elastic constant, such that the relaxation time follows a power-law distribution. (b) The same as Fig. 1(a), but we replace the white noise with the colored noise *ξ*(*t*) generated by the general Voigt model. (c-e) The same as Fig. 2(a-c) but for colored noise. (f) As we decrease the minimum relaxation time *τ*_0_ in the power-law distribution, the measured exponent approaches the predicted value 0.3. In (a-e), we set the dimensionless noise strength *A*_*p*_ ≈ 6.1 × 10^−3^, *τ*_0_ = 0.01, the deterministic growth rate *a* = 0.026, and *N* = 10^4^.

Thus, the constitutive equation of each Voigt element becomes

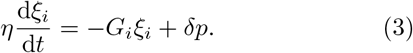

Here, *η* is the viscosity, and *G*_*i*_ is the elastic modulus of element *i*. From Eq. (3), we introduce the relaxation time as *τ*_*i*_ = *η/G*_*i*_. In the following, we non-dimensionalize the model by setting the time unit *t*_0_ = 1 min, the stress unit as *η/t*_0_, and all variables are dimensionless unless otherwise mentioned. The relaxation time is random across Voigt elements and follows a power-law distribution (Fig. 3(a)):

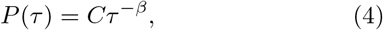

with a lower bound *τ*_0_ and the normalization factor 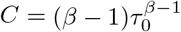

Given the overall change Δ*x* is the mean of all the Voigt elements, Δ*x* = Σ_*i*_ Δ*x*_*i*_*/N*, we obtain the modified model of cell volume growth with a colored noise:

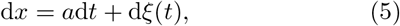

where *ξ*(*t*) = Σ_*i*_ *ξ*_*i*_(*t*)*/N* . Solving Eq. (3) for a specific element yields

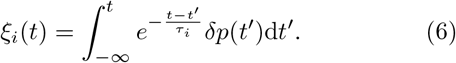

Taking account of the randomly distributed relaxation time, the overall cell growth noise *ξ*(*t*) becomes

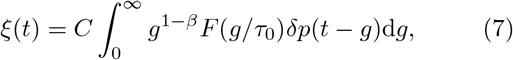

where 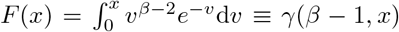 is the lower incomplete gamma function. Given Eq. (7), we find the mean square displacement of *ξ*(*t*) as

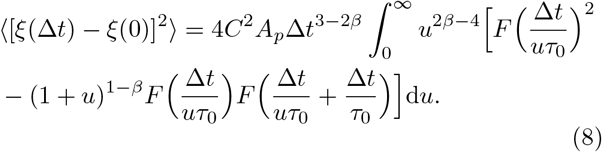

Intriguingly, the integral in Eq. (8) approaches a constant in the limit Δ*t/τ*_0_ 1 given ≫ 1 *< β <* 3*/*2, which yields the analytical relation *α* = 3 − 2*β*; consistently, we infer *β* ≈ 1.37 from the measured *α* exponent (Fig. 2(f)).

To confirm whether the general Voigt model can recapitulate the experimental phenomena, we simulate the minimal model Eq. (5) with cell cycles included (Fig. 3(b)), similar to the simulations in Fig. 1. We explicitly generate the noise *ξ*(*t*) by simulating *N* = 10^4^ Voigt elements, whose relaxation times are sampled from Eq. (4) with *τ*_0_ = 0.01. We set *β* = 1.35 so that the predicted subdiffusive exponent for the MSD of logarithmic volume is *α* = 0.3. By averaging *ξ*_*i*_ over all elements, we obtain the time-dependent noise *ξ*(*t*) in Eq. (5) (End Matter). Applying the same data analysis protocol as in Fig. 2, we find that the minimal model combined with the general Voigt model successfully recapitulates the subdiffusive random growth of bacterial cell volume (Fig. 3(c-e)).

To verify whether the measured exponent approaches the predicted value in the limit of Δ*t/τ*_0_ ≫ 1, we change *τ*_0_ (with *A*_*p*_ and *N* changing correspondingly to ensure the prefactor of the long-term subdiffusion and the expected maximum relaxation time remain invariant). Indeed, the measured exponent approaches the predicted value *α* = 0.3 as *τ*_0_ decreases (Fig. 3(f)). We also simulate an extended model in which the deterministic growth rate in each cell cycle is random, with the noise intensity chosen to match the experimental data; we confirm that the subdiffusive scaling is insensitive to fluctuations in the deterministic growth rate (Fig. S7).

## Discussion

In this work, by employing a minimal drifted Brownian motion model as a controlled benchmark, we first identify inherent statistical biases in growth rate measurements. This analysis confirms that the superexponential growth observed in the late cell cycle is a genuine biological feature rather than a sampling artifact.

More importantly, we demonstrate that the stochastic growth of bacterial cell volume exhibits subdiffusive fluctuations, a phenomenon that cannot be accounted for by white-noise growth models. The subdiffusion with an anomalous exponent *α* ≈ 0.27 suggests that negative temporal correlations govern growth fluctuations. This “memory” in the growth process occurs on the scale of minutes, which is significantly shorter than the time scale of growth rate noise generated by biochemical processes—such as gene expression, protein synthesis, or cellular metabolism—that operate on the scale of hours or a doubling time [11]. Instead, our results point toward a mechanical origin. The success of the general Voigt model, treating the cell wall as a complex viscoelastic network with a power-law distribution of relaxation times, suggests that the peptidoglycan meshwork is not merely a plastic shell. Instead, it is a complex material whose multi-scale relaxation dynamics fundamentally constrain the stochastic nature of cell growth.

The broad distribution of relaxation times implies significant heterogeneity in the cell wall’s mechanical properties, potentially stemming from stochastic bond cleavage and glycan strand insertion. Our finding relates the exponent *α* governing macroscopic growth statistics to the exponent *β* governing microscopic material parameters. Future work integrating these mechanical fluctuations with the biochemical signaling pathways of cell division may reveal how bacteria harmonize physical constraints with regulatory programs to maintain stable growth.

We note that the statistical bias in the average growth rate also persists for more complex models, where the time-averaged growth rate of each cell cycle is random, or the noise is colored (Figs. S3, S7). Beyond bacterial growth, our study highlights the need for rigorous null models when analyzing biological data. The fact that a minimal model with a strictly constant growth rate and Brownian growth-rate noise can exhibit apparent growth acceleration serves as a cautionary note.

We thank Poyi Ho and Ziming Zhao for helpful discussion related to this work. The research was funded by the National Key Research and Development Program of China (2024YFA0919600) and supported by Peking-Tsinghua Center for Life Sciences grants.

## End Matter

### Details of numerical simulations

We perform numerical simulations of the stochastic growth dynamics using a multi-scale integration scheme. The stochastic differential equation (SDE) for cell volume growth [Eq. (1)] is integrated with a time step of 0.05 min. To match the 1-min sampling interval of the experimental data [5], we sample the simulation trajectories every 20th data point. The division process follows the adder model, with noise magnitudes in both the SDE and the division volume tuned to recapitulate the observed volume trajectories in experiments.

For the viscoelastic growth model (Fig. 3), we generate the colored noise *ξ*(*t*) by simulating *N* = 10^4^ Voigt elements with a time step of 0.001 min when *τ*_0_ = 0.01. The SDE for cell volume growth [Eq. (5)] is integrated with a time step of 0.05 min. To ensure the colored noise statistics reach a stationary state, we implement a “pre-warming” period for each lineage in which the Voigt model is equilibrated for 10 min before being coupled to the volume SDE. The colored noise *ξ*(*t*) is generated independently for each simulated lineage.

Note that given *N* Voigt elements, the maximum relaxation time scales as *τ*_max_ ∼ *τ*_0_*N* ^1*/*(*β*−1)^. In Fig. 3(f), we change *τ*_0_, and in the meantime, change *A*_*p*_ as *A*_*p*_(*τ*_0_) = *A*_*p*_(*τ*_0_ = 0.01)(*τ*_0_*/*0.01)^2−2*β*^ and *N* (*τ*_0_) = *N* (*τ*_0_ = 0.01)(*τ*_0_*/*0.01)^1−*β*^ to ensure the prefactor of the MSD (Eq. (8)) and the maximum relaxation time remain invariant.

### Mechanisms of the statistical biases in the cell cycle-dependent growth rate

The averages in Fig. 1(c-h) are over bins based on the corresponding x-axes; for the average conditioned on the relative age, the weight of each data point is inversely proportional to the number of points in its corresponding cell cycle.

To understand the acceleration in the average growth rate in relative-age statistics, let us consider cells that are about to reach the division size. For these cells, their instantaneous growth rates are dominated by the random term in Eq. (1); thus, the average growth rate over these cells should scale as *λ* ∼ *δt*^−1*/*2^ where *δt* is the time left before division. Since this argument does not rely on the doubling time, the average growth rate conditioned on the relative age *ϕ* should also diverge as (1 − *ϕ*)^−1*/*2^ (Fig. S2). According to this argument, the growth rate divergence should be invariant of the time interval Δ*t* used to compute the average growth rate, which we confirm numerically (Fig. S3(a)). We provide a detailed analytical derivation strictly for the drifted Brownian model in the Supplemental Material.

We speculate that a similar statistical bias holds for colored noise, such as that generated by the general Voigt model. Specifically, if the MSD of the logarithmic volume scales as 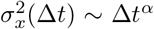, we expect the apparent growth rate to diverge as *λ* ∼ (1 − *ϕ*)^*α/*2−1^ near the cell-cycle end. Nevertheless, we find it is numerically challenging to measure the divergence exponent from simulations accurately.

For the relative-volume statistics, the reduction of the growth rate near the end of the cell cycle arises because a cell whose instantaneous growth rate is high at the end of the cell cycle can miss the last ln 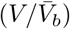 bin in its cell-volume trajectory. Therefore, the average growth rate is biased toward cells with slower instantaneous growth rates. Because this sampling bias depends on the bin size, the bias should diminish as we decrease the time interval Δ*t* to compute the growth rate, in agreement with our numerical simulations (Fig. S3(b)). This same mechanism accounts for the growth rate reduction observed in the elapsed-time statistics (Fig. S3(c)).

