## Supplemental Material for "Subdiffusive random growth of bacteria"

Jialiang Wei,[1] Yichen Yan,[2,3] and Jie Lin[2,3,*]
[1]School of Physics, Peking University, Beijing, China
[2]Center for Quantitative Biology, Peking University, Beijing, China
[3]Peking-Tsinghua Center for Life Sciences, Peking University, Beijing, China
(Dated: April 9, 2026)

### A. Analytical calculation of the bias of the average growth rate near the end of cell cycle

As shown in the main text, the "Relative Age" statistics applied to cell growth trajectories reveal an apparent acceleration in the final stage of the cell cycle (Fig. 1(c)). To understand this apparent measurement bias let us consider a simpler model where the birth volume $V_b = 1$ and the division volume $V_d = 2$, which simplifies the derivations without affecting the growth rate divergence near the cell-cycle end.

We first consider all cell-volume trajectories whose cell volume is precisely at the division threshold $V_d$ at $t = 26.7$ min. Since the cell volume dynamics include both a deterministic growth term and a random term (Eq. (1) in the main text), the cell volume can decrease due to random fluctuations, as observed experimentally. We plot multiple trajectories satisfying the condition in Fig. S1, and notice that in some of the trajectories (e.g., the blue one), the cell volume already crosses the threshold before $t = 26.7$min. These trajectories correspond to cells that divide earlier, and their actual doubling times are shorter than 26.7 min. Therefore, to compute the average growth rate as a function of the relative age for all cells that divide at 26.7 min, we have to get rid of those trajectories that divide earlier. Interestingly, the remaining trajectories tend to have higher growth rates before $t = 26.7$ min, so the average growth rate based on them is higher. If this picture holds for cells with a doubling time of 26.7 min, it should hold for other doubling times as well. Therefore, the average growth rate conditioned on relative age, regardless of doubling time, should exhibit the same bias.

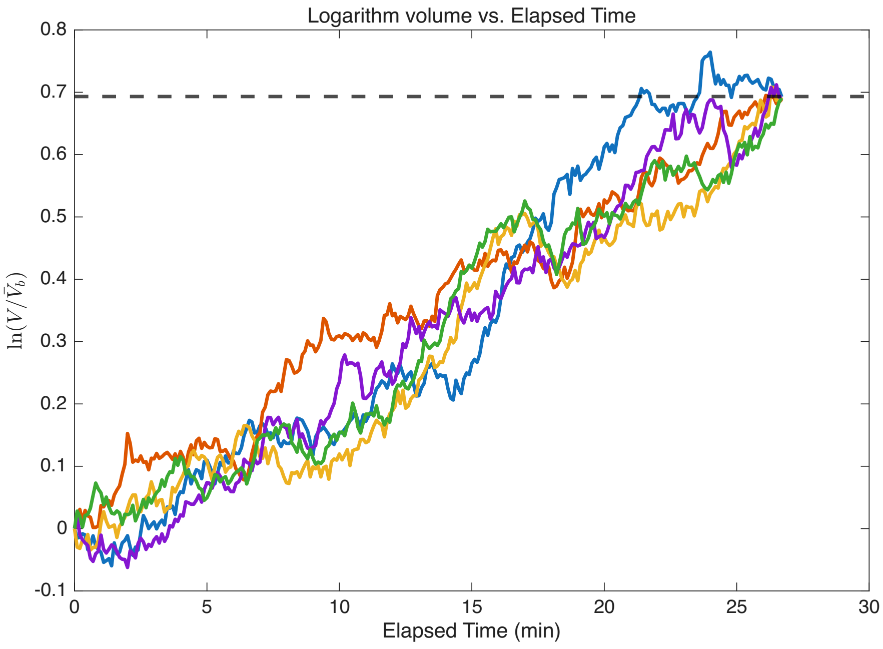


FIG. S1. Five logarithmic-volume trajectories starting at 0. The figure shows that the blue trajectory reaches the division threshold ($\ln 2$) prematurely, so it is not counted when we calculate the average growth rate for cells with a doubling time of 26.7 min.

In the following, we present a detailed mathematical derivation for the average growth rate in the relative-age statistics. Mathematically, the doubling time distribution can be modelled as a first-passage time problem of the biased Brownian motion:

$$\mathrm{d}x = a\mathrm{d}t + b\mathrm{d}W(t). \tag{S1}$$

Cell growth involves a logarithmic volume $x$ starting at $x_0 = 0$ at $t = 0$, which stops (cell division occurs) when the logarithmic volume reaches the threshold $x_{\text{div}} = \ln 2$. We introduce the relative age $\phi \in [0, 1]$ as the ratio of elapsed time $t$ to the doubling time $T$:

$$\phi = \frac{t}{T}. \tag{S2}$$

The curve in Fig. 1(c) in the main text is obtained by computing the average growth rate $\lambda(\phi)$ over all cell cycles conditioned on the relative age $\phi$. To find $\lambda(\phi)$, we decompose it into two subproblems:

1. Doubling time distribution $P(T)$: Derive the probability distribution of doubling time $P(T)$ for the biased Brownian motion, Eq. (S1).

2. Conditional average growth rate $\lambda(\phi|T)$: Determine the average growth rate for all cell cycles that share the same doubling time $T$.

$\lambda(\phi)$ is then obtained as

$$\lambda(\phi) = \int_0^\infty \lambda(\phi|T)P(T)\mathrm{d}T. \tag{S3}$$

*Doubling time distribution*

We define the survival function $S(x, t)$ as the probability that the cell has not yet divided (i.e., the logarithmic volume has not reached the boundary $\ln 2$) at time $t$, given that it started at $x$ at time $t = 0$:

$$S(x, t) = \int_{-\infty}^{\ln 2} p(x', t|x, 0)\mathrm{d}x', \tag{S4}$$

where $p(x', t|x, 0)$ is the probability density function of $x'$ at time $t$ with the boundary condition $p(\ln 2, t|x, 0) = 0$. Therefore, the initial condition and boundary condition for $S(x, t)$ are:

$$S(x, 0) = \begin{cases} 1, & \text{if } x < \ln 2 \\ 0, & \text{if } x \geq \ln 2 \end{cases} \tag{S5}$$

and

$$S(\ln 2, t) = 0. \tag{S6}$$

We note that the probability density function $p(x', t'|x, t)$ satisfies the backward Fokker-Planck equation (BFPE):

$$\frac{\partial}{\partial t}p(x', t'|x, t) = -a\frac{\partial}{\partial x}p(x', t'|x, t) - \frac{1}{2}b^2\frac{\partial^2}{\partial x^2}p(x', t'|x, t). \tag{S7}$$

Using time-translation invariance, we can rewrite Eq. (S7) as

$$\frac{\partial}{\partial t}p(x', t|x, 0) = a\frac{\partial}{\partial x}p(x', t|x, 0) + \frac{1}{2}b^2\frac{\partial^2}{\partial x^2}p(x', t|x, 0). \tag{S8}$$

Integrating Eq. (S8) over $x$ from $-\infty$ to $\ln 2$ yields the governing PDE for $S(x, t)$, subject to the absorbing boundary condition $S(\ln 2, t) = 0$:

$$\frac{\partial}{\partial t}S(x, t) = a\frac{\partial}{\partial x}S(x, t) + \frac{1}{2}b^2\frac{\partial^2}{\partial x^2}S(x, t). \tag{S9}$$

We introduce the Green's function $G(x, t|x', 0)$ of Eq. (S9), which is the solution of Eq. (S9) given $S(x, 0) = \delta(x - x')$. By introducing an imaginary particle in the space of $x > \ln 2$, we find the analytical expression of the Green's function:

$$G(x,t|x',0)=\frac{1}{\sqrt{2\pi b^2 t}}\left(e^{-\frac{(x-x'+at)^2}{2b^2t}}-e^{-\frac{2a(\ln 2-x')}{b^2}}e^{-\frac{(x-2\ln 2+x'+at)^2}{2b^2t}}\right). \tag{S10}$$

$S(x,t)$ is then found by integrating the initial condition over the Green's function:

$$S(x,t)=\int_{-\infty}^{\ln 2}G(x,t|x',0)\mathrm{d}x'. \tag{S11}$$

By integrating the initial condition (assuming the starting point $x_0$ is integrated over the Green's function, or by setting $x'=x_0$ and integrating over $x_0$), the analytical expression for $S(x,t)$ is derived using the Error Function (erf) expression:

$$S(x,t)=\frac{1}{2}\left[1+\mathrm{erf}\left(\frac{\ln 2-x-at}{\sqrt{2b^2t}}\right)-e^{\frac{2a(\ln 2-x)}{b^2}}\left(1+\mathrm{erf}\left(\frac{x-\ln 2-at}{\sqrt{2b^2t}}\right)\right)\right]. \tag{S12}$$

Setting the starting logarithmic volume $x=0$ (i.e., $x_0=0$), the expression simplifies to $S(t)$:

$$S(t)=\frac{1}{2}\left[1+\mathrm{erf}\left(\frac{\ln 2-at}{\sqrt{2b^2t}}\right)-2^{\frac{2a}{b^2}}\left(1-\mathrm{erf}\left(\frac{\ln 2+at}{\sqrt{2b^2t}}\right)\right)\right]. \tag{S13}$$

The probability density function $P(T)$ of doubling time is given by $P(T)=-\frac{\mathrm{d}S(T)}{\mathrm{d}T}$, which yields

$$P(T)=\frac{\ln 2}{\sqrt{2\pi b^2T^3}}e^{-\frac{(\ln 2-aT)^2}{2b^2T}}. \tag{S14}$$

*Conditional average growth rate*

We introduce $p_1(x,t)$ as the probability density distribution of all trajectories starting from $t=0, x=0$ and remaining below the division threshold $\ln 2$ up to time $t$. $p_1(x,t)$ satisfies the forward Fokker-Planck equation (FFPE):

$$\frac{\partial}{\partial t}p_1(x,t)=-a\frac{\partial}{\partial x}(p_1(x,t))+\frac{1}{2}b^2\frac{\partial^2}{\partial x^2}(p_1(x,t)),$$

with the initial condition $p_1(x,0)=\delta(x)$ and the absorbing boundary condition $p_1(\ln 2,t)=0$. We can find the solution for $p_1(x,t)$ as

$$p_1(x,t)=\frac{1}{\sqrt{2\pi b^2t}}\left(e^{-\frac{(x-at)^2}{2b^2t}}-e^{-\frac{(x-2\ln 2-at)^2}{2b^2t}}e^{\frac{2\ln 2a}{b^2}}\right). \tag{S15}$$

We next introduce $p_2(x,t)$ as the probability density distribution of all trajectories ending exactly at $t=T, x=\ln 2$ and remaining below $\ln 2$ at all times $t<T$. $p_2(x,t)$ satisfies the backward Fokker-Planck Equation:

$$\frac{\partial}{\partial t}p_2(x,t)=-a\frac{\partial}{\partial x}p_2(x,t)-\frac{1}{2}b^2\frac{\partial^2}{\partial x^2}p_2(x,t),$$

with the initial condition set at the final state $p_2(x,T)=\delta(x-\ln 2)$ and the boundary condition $p_2(\ln 2,t)=0$. We find that $p_2(x,t)$ has the following property:

$$p_2(x,t)\propto\frac{1}{\sqrt{2\pi b^2(T-t)}}\frac{\ln 2-x}{T-t}e^{-\frac{(\ln 2-x-a(T-t))^2}{2b^2(T-t)}}. \tag{S16}$$

We next introduce the probability density function $p(x,t|T)$ of all trajectories that start at $x=0$ at $t=0$ and reach the division threshold $x=\ln 2$ the first time at $t=T$, which is related to $p_1$ and $p_2$ by

$$p(x,t|T)\propto p_1(x,t)p_2(x,t). \tag{S17}$$

Obviously, when $x\geq\ln 2$, $p(x,t|T)=0$. We next compute the average position $x(t)$ at time $t$ over $p(x,t|T)$

$$x(t|T)=\frac{\int_{-\infty}^{\ln 2}xp(x,t|T)\mathrm{d}x}{\int_{-\infty}^{\ln 2}p(x,t|T)\mathrm{d}x}, \tag{S18}$$

where $p(x,t|T)$ is the conditional probability density function.

The average normalized growth rate trajectory, $\lambda(t|T)$, is obtained by differentiating the conditional expected position, $x(t|T)$, where the expected position is calculated through the spatial integration of the logarithmic volume $x$ against the conditional probability density $p(x,t|T)$. Finally, $\lambda(t|T)$ is normalized to the time scale $\phi \in [0,1]$ to yield $\lambda(\phi|T)$.

The analytical solution for $x(t|T)$ turns out to be

$$x(t|T) = \ln 2 - \left[\left((1-\frac{t}{T})\ln 2 + \frac{b^2 t}{\ln 2}\right)\operatorname{erf}\left(\ln 2\sqrt{\frac{T-t}{2b^2 tT}}\right) + \sqrt{\frac{2b^2(T-t)t}{\pi T}}e^{-\frac{(\ln 2)^2(T-t)}{2b^2 Tt}}\right]. \tag{S19}$$

The conditional average growth rate $\lambda(t|T)$ is the time derivative of the average position: $\lambda(t|T) = \frac{\mathrm{d}x(t|T)}{\mathrm{d}t}$, which is

$$\lambda(t|T) = \left(\frac{\ln 2}{T} - \frac{b^2}{\ln 2}\right)\operatorname{erf}\left(\ln 2\sqrt{\frac{T-t}{2b^2 tT}}\right) + \sqrt{\frac{2tb^2}{\pi T(T-t)}}e^{-\frac{(\ln 2)^2(T-t)}{2b^2 Tt}}. \tag{S20}$$

We next convert the conditional average growth rate $\lambda(t|T)$ as a function of the relative age $\phi \in [0,1]$ by substituting $t = \phi T$:

$$\lambda(\phi|T) = \left(\frac{\ln 2}{T} - \frac{b^2}{\ln 2}\right)\operatorname{erf}\left(\ln 2\sqrt{\frac{1-\phi}{2b^2 \phi T}}\right) + \sqrt{\frac{2\phi b^2}{\pi T(1-\phi)}}e^{-\frac{(\ln 2)^2(1-\phi)}{2b^2 \phi T}}. \tag{S21}$$

The average growth rate $\lambda(\phi)$ over all cell cycles conditioned on the relative age $\phi$ is obtained by integrating the conditional average growth rate $\lambda(\phi|T)$ with the doubling time distribution $P(T)$ over $T$:

$$\lambda(\phi) = \int_0^{+\infty} \frac{\ln 2}{\sqrt{2\pi b^2 T^3}}e^{-\frac{(\ln 2 - aT)^2}{2b^2 T}}\left[\left(\frac{\ln 2}{T} - \frac{b^2}{\ln 2}\right)\operatorname{erf}\left(\ln 2\sqrt{\frac{1-\phi}{2b^2 \phi T}}\right) + \sqrt{\frac{2\phi b^2}{\pi T(1-\phi)}}e^{-\frac{(\ln 2)^2(1-\phi)}{2b^2 \phi T}}\right]\mathrm{d}T. \tag{S22}$$

We confirm the validity of Eq. (S22) by comparing it with direct numerical simulations similar to Fig. 1 in the main text except the birth and division volumes are fixed at $V_b = 1$ and $V_d = 2$ (Fig. S2).

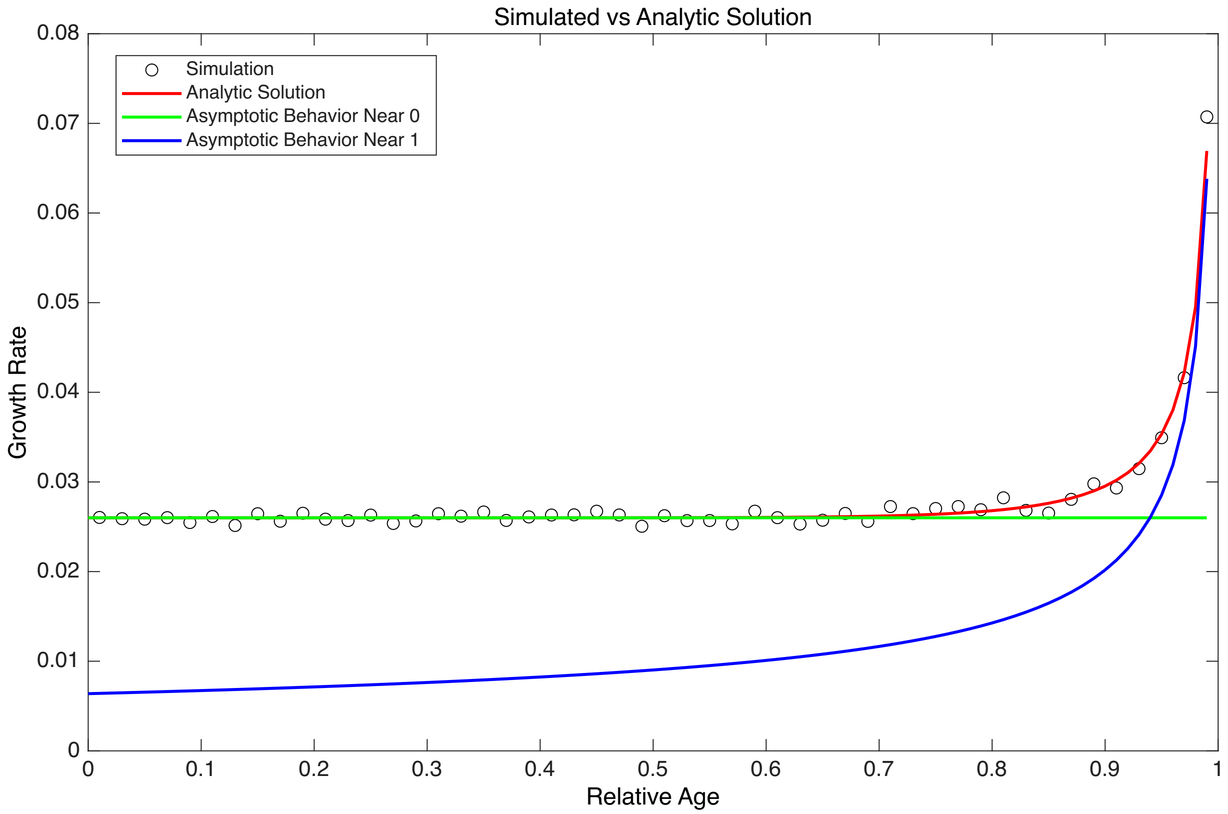


FIG. S2. The theoretical prediction Eq. (S22) agrees with the numerical simulation. We also show the predicted asymptotic behaviors near $\phi \to 0$ and $\phi \to 1$.

In the limit where the relative age $\phi \to 0$, the average growth rate approaches the deterministic growth rate $a$:

$$\lim_{\phi \to 0} \lambda(\phi) \approx \int_0^{+\infty} \frac{\ln 2}{\sqrt{2\pi b^2 T^3}} e^{-\frac{(\ln 2 - aT)^2}{2b^2 T}} \left( \frac{\ln 2}{T} - \frac{b^2}{\ln 2} \right) \mathrm{d}T = a. \tag{S23}$$

In the limit where $\phi \to 1$, the error function term tends to zero and the average growth rate turns out to be

$$\lim_{\phi \to 1} \lambda(\phi) \approx \frac{2a}{\pi\sqrt{1-\phi}} 2^{a/b^2} K_1 \left( \frac{a \ln 2}{b^2} \right), \tag{S24}$$

where $K_1(x)$ is the modified Bessel function of the second kind of order one. This analysis reveals a critical finding: as $\phi \to 1$, the average normalized growth rate $\lambda(\phi)$ diverges with the asymptotic form:

$$\lambda(\phi) \propto \frac{1}{\sqrt{1-\phi}}. \tag{S25}$$

To estimate the range where this selection bias ($\phi \to 1$ divergence) dominates, we define the boundary range by the intersection point where the asymptotic solution $\lambda(\phi)|_{\phi \to 1}$ equals $a$. By equating the two asymptotic solutions, the intersection point $1-\phi$ is found to be:

$$1 - \phi \approx \left[ \frac{2}{\pi} 2^{a/b^2} K_1 \left( \frac{a \ln 2}{b^2} \right) \right]^2 \equiv f\left( \frac{a}{b^2} \right). \tag{S26}$$

The function $f(a/b^2)$ is a monotonically decreasing function of the ratio $a/b^2$. It is clear that even if the deterministic growth rate ($a$) and the noise intensity ($b$) depend on the cell cycle, e.g., via the cell volume, the exponent of the divergence of the average growth rate near the end of the cell cycle ($\phi \to 1$) remains robustly fixed at 0.5.

*

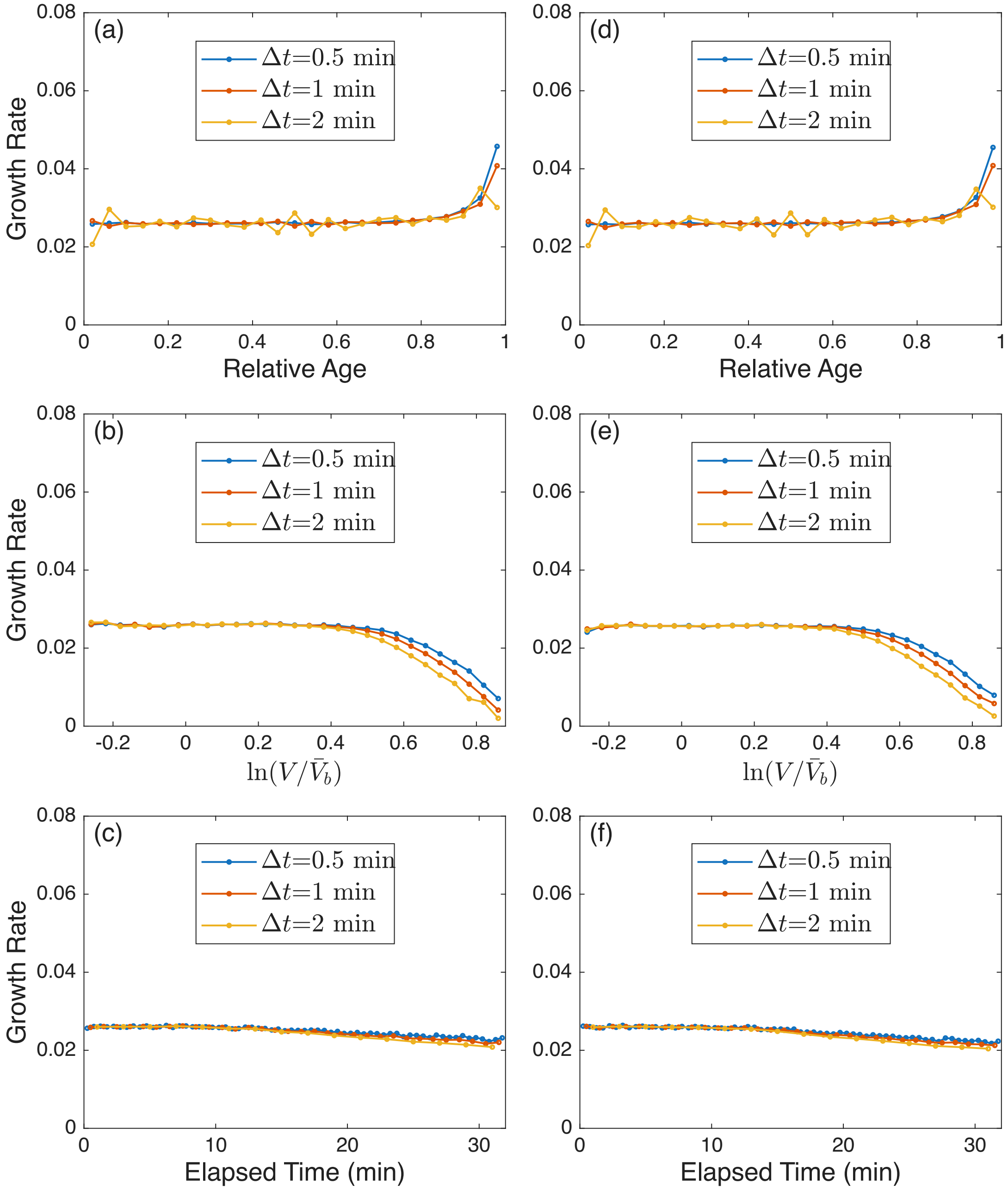


FIG. S3. Cell cycle-dependent growth rate for different time intervals and additional models. (a-c) The same analysis as Fig. 1(c-e) in the main text, but for different time intervals to compute the growth rate. For the relative-age statistics, the apparent growth acceleration near the cell-cycle end does not diminish as $\Delta t$ decreases. In contrast, for the relative-volume and elapsed-time statistics, the deviation from the ground-truth constant growth rate (a straight horizontal line) decreases with a smaller $\Delta t$, confirming that this reduction bias stems from the finite time interval. (d-f) The same analysis as (a-c), but the deterministic growth rate varies across cell cycles with a coefficient of variation (CV) of 0.1, consistent with experimental measurements [1]. The statistical bias is robust against the randomness in the deterministic growth rate across cell cycles.

[1] S. Taheri-Araghi, S. Bradde, J. T. Sauls, N. S. Hill, P. A. Levin, J. Paulsson, M. Vergassola, and S. Jun, Cell-size control and homeostasis in bacteria, Current Biology **25**, 385 (2015).

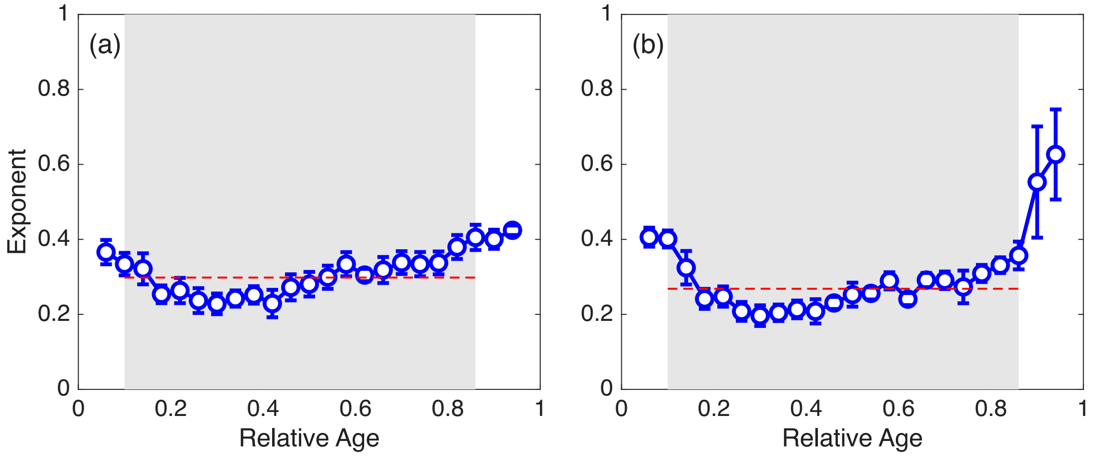


FIG. S4. The same analysis as Fig. 2(f) in the main text, but for the data of cell length (a) and surface area (b).

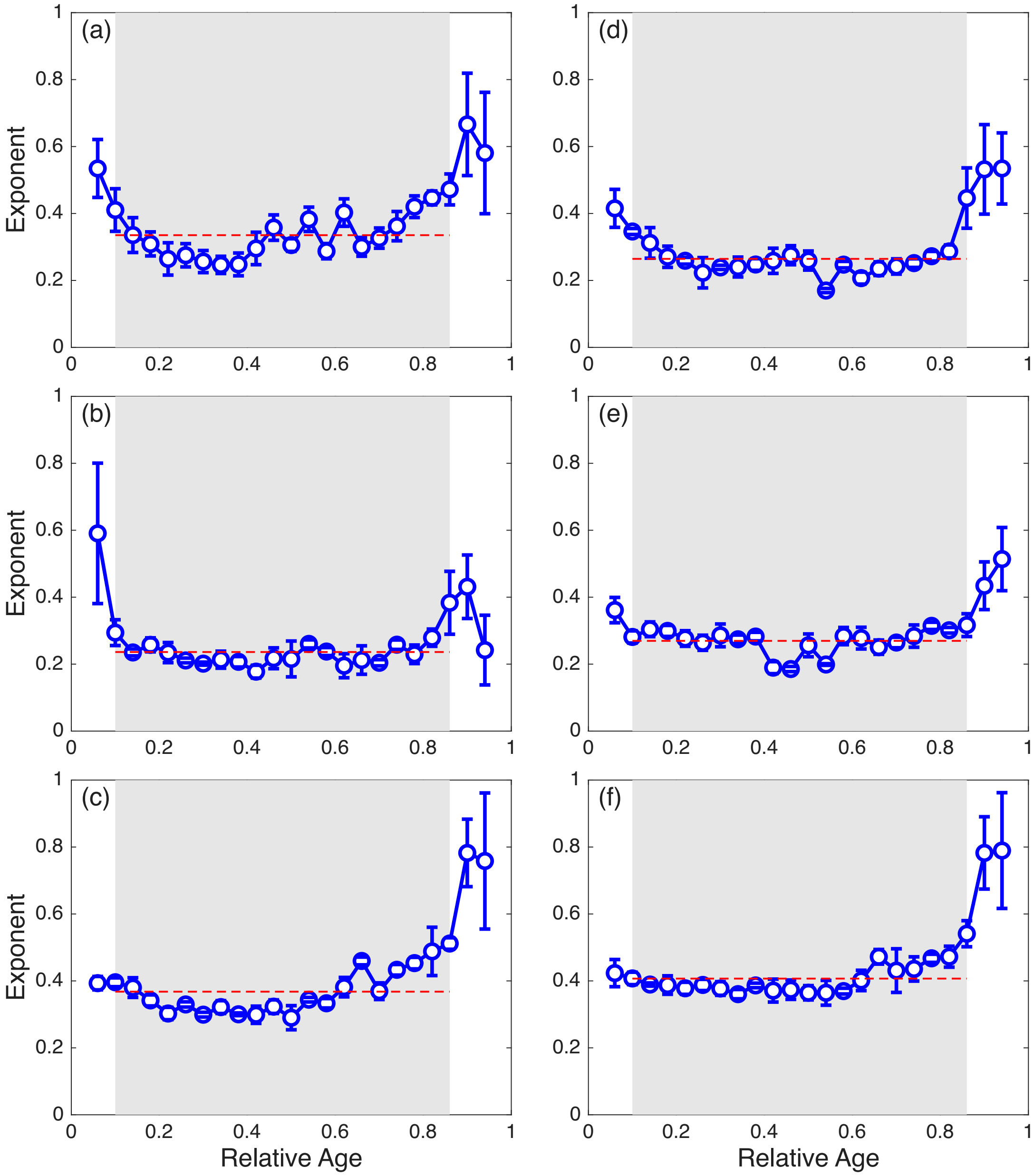


FIG. S5. The same analysis as Fig. 2(f) in the main text, but for different growth mediums, including (a) glucose + 6 a.a (b) glucose, (c) glycerol, (d) sorbitol, (e) synthetic rich, and (f) tryptic soy broth (TSB) according to Ref. [1].

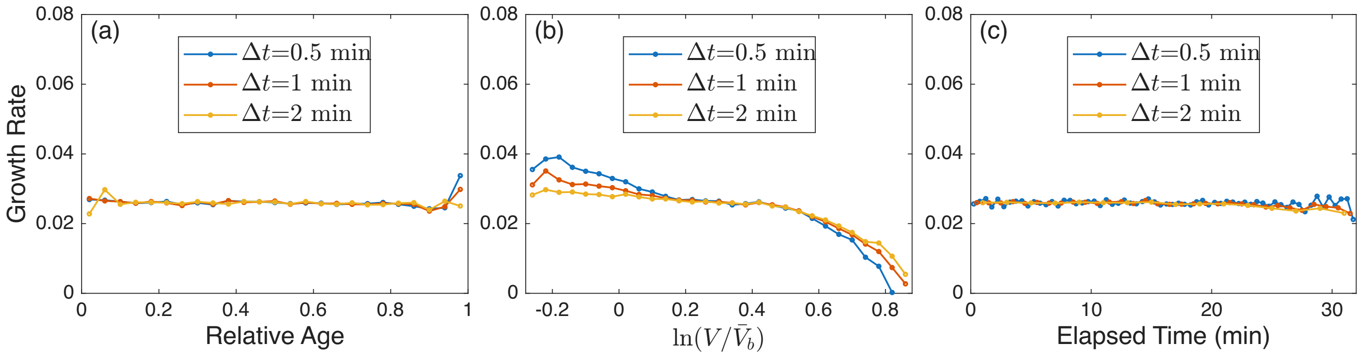


FIG. S6. The same analysis as Fig. S3(a-c), but the growth rate noise is generated from the general Voigt model with $\beta = 1.35$. We note that for the relative-volume statistics, the deviation from the ground truth is more significant as $\Delta t$ decreases.

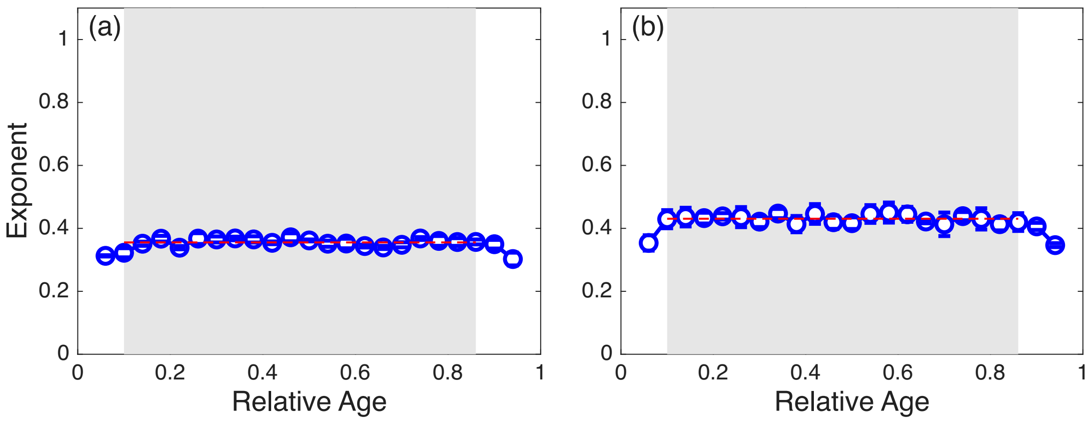


FIG. S7. The same analysis as in Fig. 3(e) of the main text, but for the extended model in which the deterministic growth rate is random across cell cycles. We sample the deterministic growth rate for each cell cycle from a normal distribution with mean equal to 0.026 and the coefficient of variation equal to 0.1 in (a) and 0.2 in (b), which are biologically reasonable values [1]. We find that the subdiffusive random growth remains robust.
